# *De novo* non-compartmentalized microorganism for directed evolution of nanobodies

**DOI:** 10.1101/2024.11.27.625622

**Authors:** Donglian Wu, Huiping He, Ruyu Xi, Simin Xia, Xi Chen

## Abstract

The creation of *de novo* microorganisms is a fascinating task, and microorganisms offer a wide range of applications for humanity. Herein, we designed a *de novo* microorganism, termed Syn-phage, as an artificial unencapsulated mimic of naturally occurring phage capable of evolving nanobodies with improved properties. Syn-phage composes unified genomic and phenotypic materials with phenotypic proteins encoded from its genetic information. Since Syn-phage can undergo amplification in *E. coli*. host, we established a protocol for the evolution of nanobodies with improved avidity or affinity, allowing super-resolution live-cell immunoimaging of endogenous proteins, a challenge posed in immunolabeling with antibodies. Utilizing a Syn-phage evolved nanobody, we visualized the microtubule nucleation factor hTPX2, recorded the dynamics of it during mitotic progression, and imaged it at a resolution beyond the diffraction limit of light via an AI-powered imaging pipeline. Hence, this study demonstrated the possibility to design *de novo* microorganisms for capitative applications.

## Introduction

The creation of *de novo* microorganism is a captivating endeavor within the fields of synthetic biology and artificial life design^1, 2^. Scientists have successfully engineered a bacterial cell with a minimal genome, serving as a versatile platform for investigating core life functions and exploring whole-genome design^3^. In addition to synthesis of prokaryotic genomes^4^, eukaryotic yeast genomes were also assembled allowing us to ask otherwise intractable questions about chromosome structure, function, and evolution^5, 6^. While top-down synthetic cells are modified from naturally existing organisms, true *de novo* design involves bottom-up strategies—assembly of functional modules and gene elements from scratch. These approaches are believed to not only shed light on fundamental questions about life, but also hold immense potential for breeding technological innovations^7^. However, existing works, such as artificial cell resembles that mimic natural functions, lack fully encodable genetic information, preventing them from truly emulating living organisms^8^. Consequently, designing non-nature-occurring artificial microorganisms exhibiting the essential “living” features and concise composition remains an attractive task^9^.

Microorganisms, whether naturally existing or artificially created, serve diverse purposes in scientific research and biotechnology. For instance, they have been harnessed for the development of displaying techniques^10^ and the directed evolution of enzymes^11^. Directed evolution, specifically, is a powerful strategy for enhancing or altering the activity of biomolecules^12-14^. In this context, the evolution of nanobody—camelid-derived single-chain V^HH^ domain antibody — holds immense appeal. Nanobodies are garnering increasing attention due to their high stability, compact size (∼14 kDa), and the high affinity and specificities against their binding partners^15^. Their ability to be expressed intracellularly or transduced into cells^16^ in a non-endocytic manner makes nanobodies an excellent choice for live-cell immunolabeling^17, 18^. Nevertheless, super-resolution (SR) immunolabeling of endogenous proteins inside living cells using antibodies still represents a big challenge^19^. As part of our research, we aim to leverage *de novo* created microorganisms to evolve nanobodies with improved features, such as increased affinity or improved avidity, facilitating SR live-cell immunoimaging through an AI-powered pipeline.

## Results

### *De novo* design and synthesis of a microorganism—Syn-phage

Inspired by the structural composition of viruses and phages, we rationally design an artificial semi-microorganism termed as Syn-phage (**Fig. 1a-b**). Hence, a pair of DNA binding protein (DBP) and its cognate DNA sequence should be identified in advance. Specifically, we employed the DBP called MafB which exhibits low-picomolar strong affinity for the palindromic T-MARE motif TGCTGACTCAGCA (i.e. <u>M</u>af<u>B</u>-binding <u>m</u>otif, or MBM)^20^. The high-resolution crystal structure unveiled that a MafB homodimer interacts with an MBM. To further enhance the unity between DBP and DNA, we employed three MBM repeats exhibiting tripled binding avidity supposed to interact with three MafB dimers (**Fig. 1b**). The circular DNA material composes several key gene elements sequentially listed below: 3×MBM/ T7 promoter (T7p)/ lac operator (LacO)/ ribosome binding site (RBS)/ MafB/ mCherry/ His-tag (**Fig. 1c**, left). The 3×MBM is positioned before T7p so that the MBM/MafB interaction is not going to affect subsequent transcription. A mCherry tag facilitates fluorescence or immunoblot detection of Syn-phage while the His-tag at the C-terminus allows ease of nickel immobilized metal affinity chromatography (Ni IMAC) isolation of Syn-phage. Assembled from three chemically synthesized fragments (2701bp/ 1803bp/ 1914 bp) (**Fig. 1c**, middle), the circular DNA (6148 bp) gives rise to the *de novo* Syn-phage **A** (or **v1.0**) upon rescued from an *E. coli* host (**Fig. 1c**, right).

**Fig. 1.**
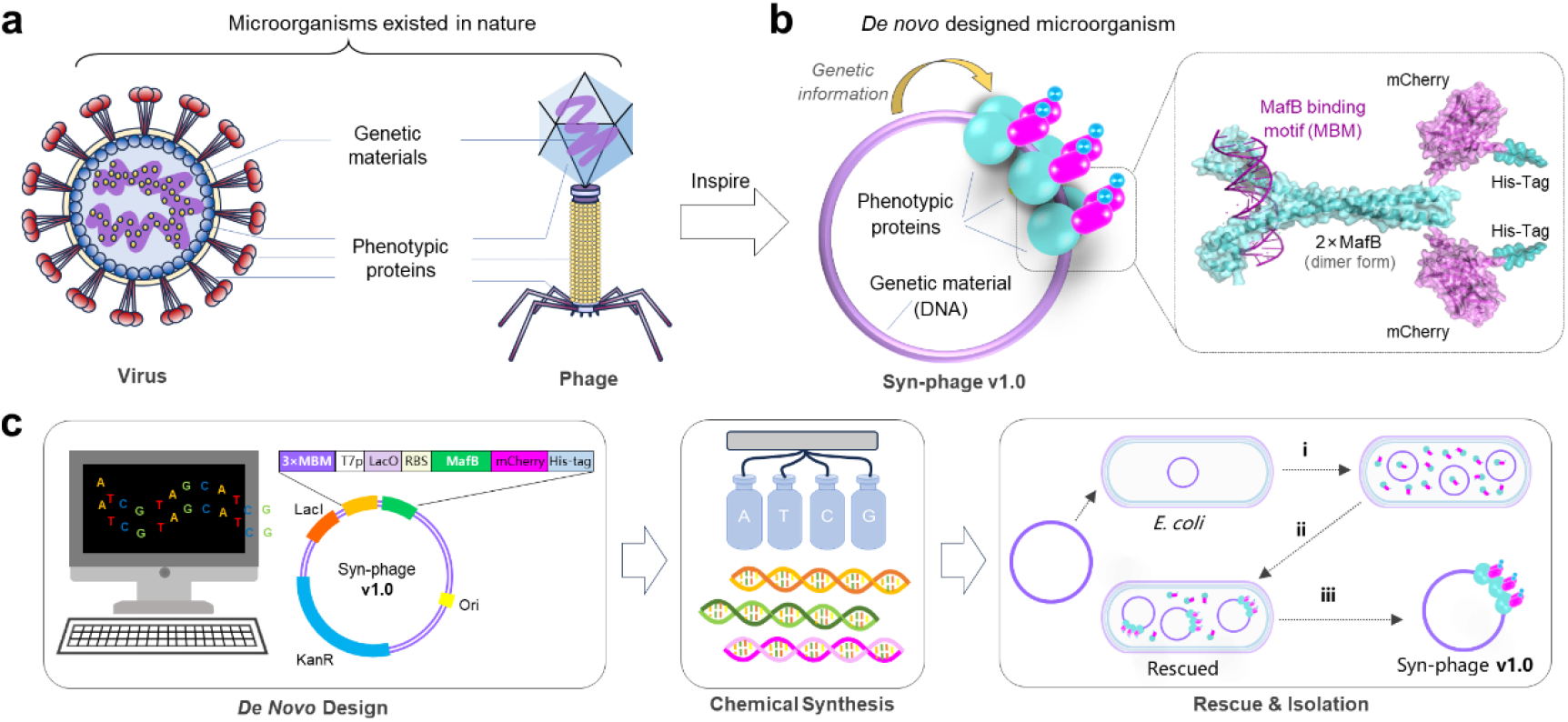
*De novo* design and creation of Syn-phage **v1.0** as an unencapsulated mimic of the nature-existing microorganism—phage. **a**, Schematic view of the structural composition of naturally existing microorganisms like virus and phage. **b**, Schematic view of the rationally designed microorganism Syn-phage which contains a circular DNA as the genetic material and six protein repeats carrying the MafB binding domain. **c**, Schematic view of *de novo* design, chemical synthesis, and host rescue for the generation of Syn-phages. In the right panel: i) DNA replication & protein expression; ii) self-assembly; iii) Ni IMAC purification.

### Characterization of Syn-phage A (i.e. v1.0)

After rescuing Syn-phage **A** from *E. coli*, we purified it through Ni IMAC followed by size-exclusion chromatography (SEC). Analytic SEC reveals a single peak with retention volume (VR) of 8.25 min (**Fig. 2a**, left). Denaturing SDS-PAGE analysis reveals a single band of around 44 kDa, matching the M.W. of the phenotypic protein material—MafB-mCherry-His^8^ (**Fig. 2a**, upper-right). The identity of this protein was further validated using Western blot (**Fig. 2a**, upper-right) and MS^2^ peptide mass fingerprinting (**Fig. 2b**). Importantly, repeated “infection” experiments confirmed that genetic information could be inherited from mother to daughter and further to the next generation (**Fig. 2c**). We also evaluated and compared Syn-phage **A** with other designed Syn-phages, including Oct1-based (Syn-phage **B**), Gal4-based (Syn-phage **C**), and MafG-based (Syn-phage **D**) variants; unfortunately, all these variants failed to efficiently infect *E. coli* host cells to produce positive colonies (**Extended Data Fig. 1**). Thus, we have successfully designed and identified a *de novo* microorganism that integrates unified genetic and phenotypic information and is capable of amplification within host cells (**Fig. 2d**).

**Fig. 2.**
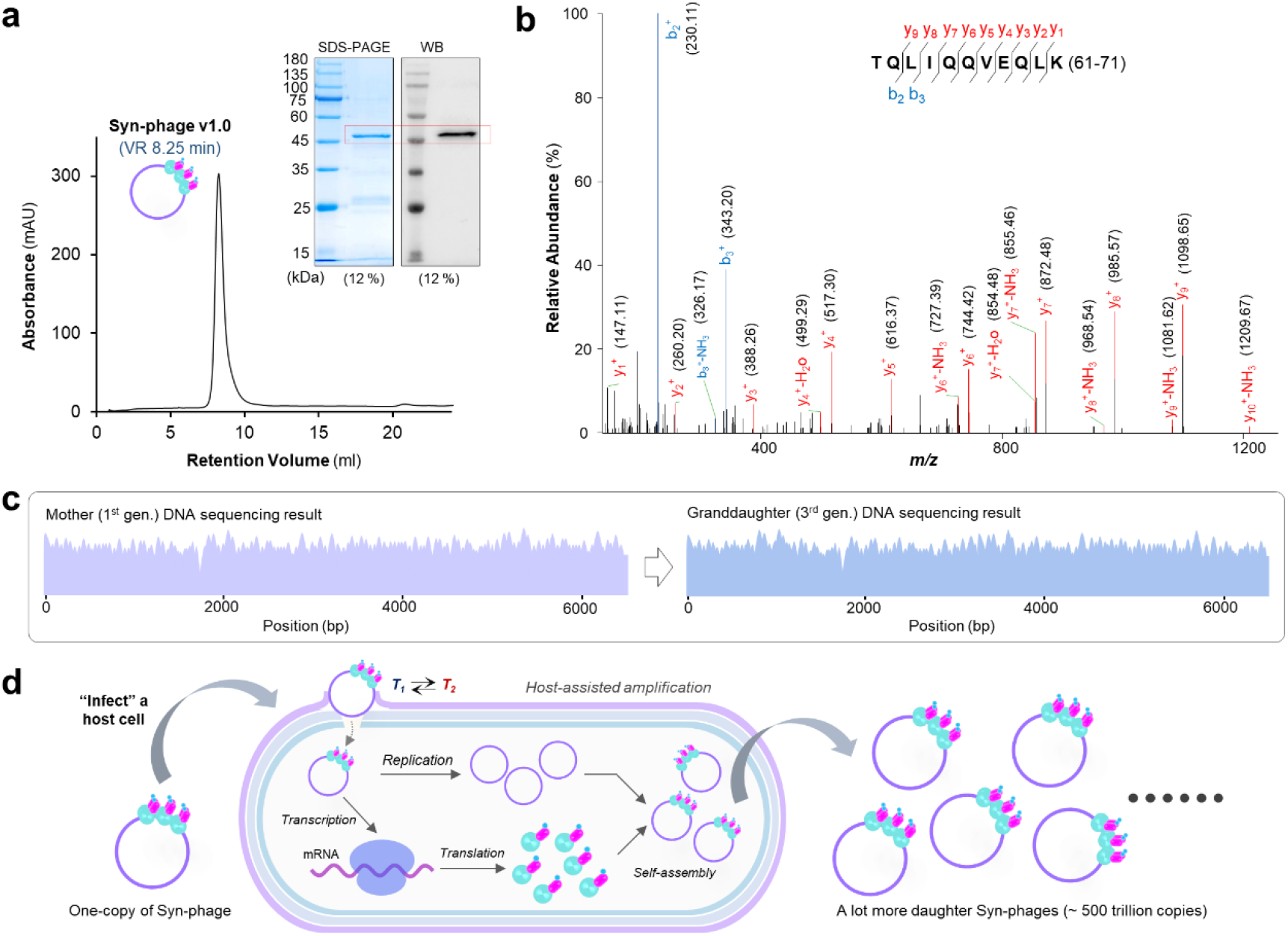
Syn-phage **v1.0** is an artificial life-like microorganism. **a**, Left: size-exclusion chromatographic (SEC) characterization of Syn-phage **v1.0** using *Superdex 200 Increase 10/300 GL* column at a flow rate of 0.4ml·min^-1^ revealing a retention volume (VR) of 8.25 ml; right: denaturing SDS-PAGE (12 %) and Western blot (anti-mCherry) analysis of Syn-phage **v1.0. b**, Tandem MS^2^ analysis of the identity of the phenotypic protein; the representative MS^2^ spectrum of TQLIQQVEQLK (61-71 of MafB) was shown. **c**, Whole genome sequencing results showed that the DNA sequence of the granddaughter (3^rd^ gen.) Syn-phage **v1.0** is identical to the parental Syn-phage **v1.0. d**, A scheme shows that a Syn-phage exhibits the essential living characteristic of a microorganism that can get amplified through “infection” of a host bacterial cell.

### Directed evolution of G6 nanobody using Syn-phage v1.1

Motivated by the above success, we next designed a protocol for evolution of nanobodies using this MafB-based Syn-phage. We were primarily interested in improving nanobodies against intrinsically disordered antigens which tend to be less or non-immunogenic during antibody generation^21^. Through screening a naïve M13 phage library (2×10^9^ pfu), we identified two lead nanobodies: G6 and F12. The two nanobodies specifically bind with the intrinsically disordered protein (IDP) called hTPX2, a crucial microtubule nucleation factor involved in cell division^22^. Additionally, we obtained a C3 nanobody by screening another naïve M13 phage library (15×10^9^ pfu) against a second IDP called BuGZ, which also plays essential roles in cell division^23^. Among these nanobodies, G6 is a very weak binder while F12 and C3 show stronger nanomolar level binding affinities.

Our designed evolution protocol comprises three key steps: (**I**) gene diversification, (**II**) rescue and selection, and (**III**) characterization of evolved nanobodies (**Extended Data Fig. 2a-b**). Initially, we conducted double site-saturated mutagenesis at two adjacent residues (R112K113, or RK) within the CDR3 region (S105-D117: SPWYFPRRKDEYD) of the G6 nanobody (**Extended Data Fig. 2c-e**). Screening the Syn-phage mutational library against TPX2-EGFP coated beads yields six promising mutants—KV, KM, PG, YG, FV and IW— due to their higher repeating frequency. Föster resonance energy transfer (FRET) assays indicated increased FRET signals, with KV performing best, closely followed by KM (**Extended Data Fig. 3a-b**). Isothermal titration calorimetry experiments revealed that the KV mutant (*K*_d_ = 8.1 μM) exhibited significantly enhanced binding affinity compared to the weak binder G6 (*K*_d_ > 100 μM) (**Extended Data Fig. 3c-e**). Live-cell colocalization assays further confirmed higher colocalization of KV and KM with hTPX2 on the mitotic spindle (**Extended Data Fig. 3f-g**). Hence, Syn-phage **v1.1** enables the evolution of a nanobody with enhanced binding affinities (**Extended Data Fig. 3h**).

### Directed Evolution of C3 against BuGZ using Syn-phage v1.2

For the evolution of C3 nanobody, we optimized the protocol by employing NNK-based single-site saturated mutagenesis across the entire CDR3 region (**Extended Data Fig. 4**). Specifically, we inserted the C3 nanobody before the MafB gene of Syn-phage **v1.0**, resulting in Syn-phage **v1.2** (**Fig. 3a-b**). Following screening against BuGZ-coated beads, we identified several promising mutants with repeat frequency of two or more. Among them, D112A, R104K, and W102A exhibited stronger binding, as revealed by live-cell colocalization assay (**Fig. 3c-d**). Notably, the W102A mutant, which displayed the highest colocalization, even facilitated the translocation of nucleus-localizing BuGZ to the mitochondria region. Further, ITC measurements revealed that the evolved nanobody W102A achieved a notable increase—from sub-nanomolar *K*_d_ of 485 nM to 81 nM with around sixfold of enhancement (**Fig. 3e-f**). Thus, our findings underscore Syn-phage’s capability to evolve nanobodies with enhanced binding affinity (**Fig. 3g**).

**Fig. 3.**
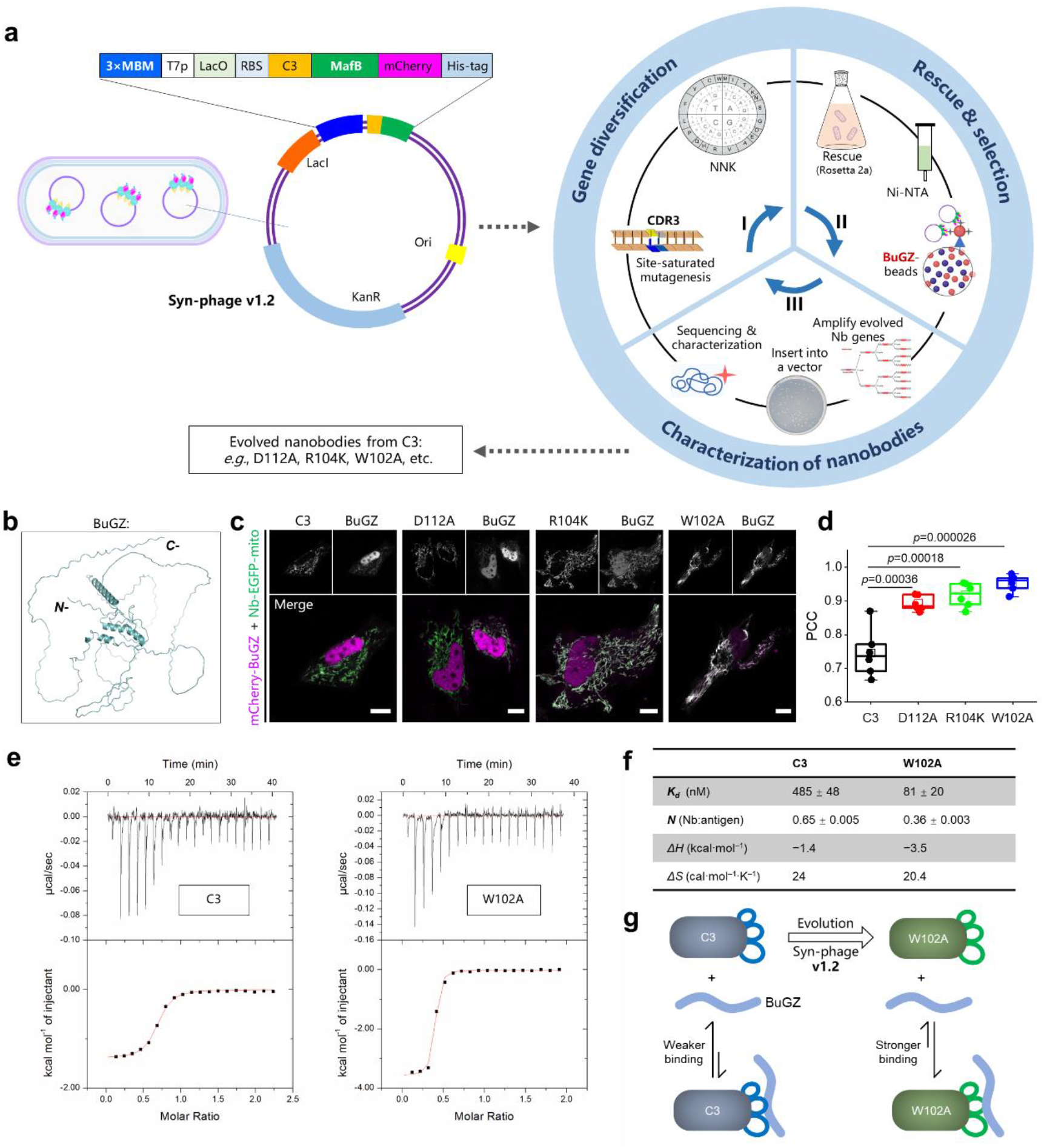
Schematic view of the evolution protocol for optimizing C3 nanobody of BuGZ. **a**, The protocol includes three major steps, namely **step I**: gene diversification, **step II**: rescue and selection, and **step III**: characterization of nanobodies. The CDR3 region of C3 was subjected to single site-saturated mutagenesis using the 32-codon trick via NNK degeneracy in **step I**; and the resultant Syn-phage **v1.2** was rescued in *E. coli*. (Rosetta 2a), purified, and selected on BuGZ-coated beads in **step II**; the nanobody gene of the selected Syn-phage **v1.2** was amplified, cloned into an appropriate vector for evaluation and the evolved nanobodies were further characterized and evaluated in **step III. b**, AlphaFold3 (https://alphafoldserver.com) predicted 3D-structure of BuGZ reveals intrinsically disordered nature of this antigen. **c**, Representative confocal micrographs of live HeLa cells co-expressing mCherry-BuGZ (magenta) and C3(mutant)-EGFP-mito (green, mitochondria) reveal that mCherry-BuGZ displays the highest colocalization with W102A-EGFP-mito. Scale bars: 10 μm. **d**, Statistical Pearson’s correlation coefficient (PCC) analysis between mCherry channel and EGFP channel reveals that the W102A mutant shows the highest colocalization degree with BuGZ (n=6 cells in each group). **e**, Isothermal titration calorimetry (ITC) results revealed that W102A mutant shows enhanced binding affinity against BuGZ than C3. **f**, A table lists the binding affinity, stoichiometry and thermodynamic parameters of the antibody-antigen binding reactions. **g**, Schematic view of the evolution of C3 to W102A resulting in enhanced binding affinity with BuGZ.

### Directed evolution of F12 against hTPX2 using Syn-phage v1.3

Next, we directed our efforts toward evolving the F12 nanobody against the intrinsically disordered and unligandable protein hTPX2. Similarly, we inserted the F12 nanobody gene before MafB, resulting in Syn-phage **v1.3** (**Fig. 4a**). We employed the same evolution protocol for C3 using single site-saturated mutagenesis across the entire CDR3 region (**Extended Data Fig. 5a-c**). Screening the mutant library against hTPX2-coated beads led us to identify four mutants with repeat frequency of two or more. Notably, two nanobody mutants—S102T and G112S—exhibit higher degree of colocalization than F12 in live-cell colocalization assay (**Extended Data Fig. 5d**). Further analysis using Pearson’s correlation coefficient (PCC) revealed that G112S-mCherry achieved a markedly enhanced PCC of over 0.9 (**Fig. 4b**). Interestingly, isothermal titration calorimetry demonstrated that the evolved G112S nanobody (*K*_d_ = 106 nM) shows only moderate enhancement in binding affinity compared to the F12 nanobody (*K*_d_ = 233 nM) (**Fig. 4c**). Therefore, we next explored another critical parameter of antibody called avidity which is closely related to binding valency^24^. From the binding thermodynamic parameters measured by ITC, we found that hTPX2 binds with F12 in a 5:1 ratio, whereas hTPX2 binds with G112S in a 1:1 ratio (**Fig. 4d**). This implies that hTPX2 could engage with approximately five times more G112S nanobody than with the F12 nanobody, resulting in around fivefold improved binding avidity (**Fig. 4e**). Consequently, the combination of enhanced binding avidity and increased affinity positions G112S as a promising candidate for biological applications, such as live-cell immunoimaging.

**Fig. 4.**
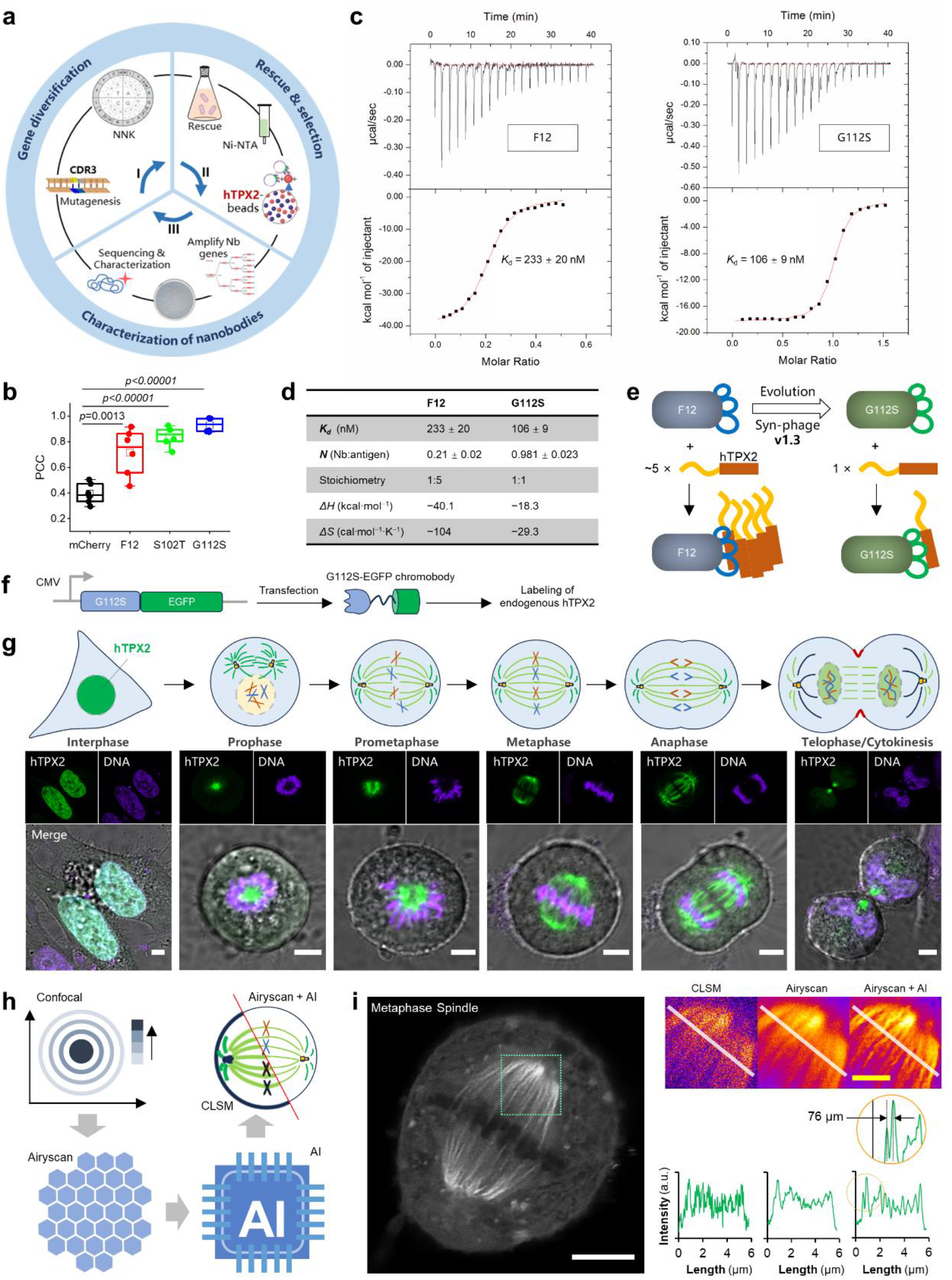
Evolution of F12 nanobody using Syn-phage **v1.3** with enhanced avidity facilitating confocal or AI-powered super-resolution (SR) live-cell immunoimaging. **a**, Schematic view of the evolution protocol; the F12 nanobody gene was inserted before MafB. **b**, Statistic quantification of the Pearson’s correlation coefficients between EGFP-hTPX2 and mCherry, F12-mCherry, S102T-mCherry, or G112S-mCherry (n=6 cells in each group). **c**, Isothermal titration calorimetry (ITC) analysis of the binding thermodynamics and stoichiometry for F12 or G112S with hTPX2 protein. **d**, A table that summarizes the dissociation constant *K*_d_, binding ratio *N*, stoichiometry, enthalpy change ΔH, and entropy change ΔS. **e**, Schematic view of the around five times of enhancement of binding avidity between nanobody and hTPX2 after the evolution of F12 to G112S. **f**, Schematic view of G112S-EGFP chromobody-based immunoimaging of endogenous hTPX2. **g**, Representative confocal images of immunolabeled hTPX2 at the interphase, prophase, prometaphase, metaphase, anaphase and telophase/cytokinesis. **h** Schematic view of the AI-assisted Airyscan super resolution immunoimaging pipeline. **i**, Left: Representative SR images of endogenous hTPX2 at metaphase spindle; right: line profile analysis reveals separation of details of less than 80 nm in distance. Scale bars: 5 μm for white, 2 μm for yellow.

### Evolved G112S nanobody enables SR immunoimaging of hTPX2

Subsequently, we proceeded to apply G112S nanobody for immunolabeling of endogenous hTPX2 using the chromobody method, which employs an expressed intrabody with a fluorescent protein (FP) tag—typically at the C-terminus—fused with an N-terminal nanobody (**Fig. 4f**). Live-cell immunolabeling of endogenous protein offers advantages over ectopically expressing FP-fused protein, which can introduce unpredictable artifacts (as observed with hTPX2, **Extended Data Fig. 6**). To our knowledge, the precise positioning of endogenous hTPX2 and its dynamics at each cell cycle stage in live mammalian cells have not been previously recorded. We observed that hTPX2 is exclusively localized within the nucleus but not on the microtubules during interphase, predominately localized to mitotic spindles from prometaphase to anaphase during M-phase, and dissociates from microtubules during cytokinesis (**Fig. 4g**). We were particularly intrigued by the rapid repositioning of hTPX2 from the mitotic spindle in anaphase to a largely non-microtubule-localized state through telophase to cytokinesis. Hence, we employed a HeLa cell line stably expressing mCherry-α-Tubulin that facilitates the visualization of microtubules and recorded a video of a dividing cell that spans from anaphase to cytokinesis. Remarkably, this acute hTPX2 translocation occurred precisely at the onset of cytokinesis, coinciding with the formation of contracting ring (**Extended Data Fig. 7a**). We propose to term this phenomenon “TPX2-escape”, signifying hTPX2’s rapid dissociation from microtubules as it repositions toward the nucleus (**Extended Data Fig. 7b**).

Finally, we employed the evolved G112S nanobody in conjunction with an AI-assisted^25^ Airyscan imaging pipeline to capture SR images of hTPX2 inside live cells (**Fig. 4h**). An Airyscan image is derived from 32 confocal images centered and surrounding each of an airy spot, resulting in an image with enhanced resolution and signal-to-noise ratio^26^. Using this imaging protocol, we observed that hTPX2 localizes in the mitotic spindle with a tendency to enrich at spindle poles, while being nearly absent in the chromosomes (**Fig. 4i**, left). By comparing confocal, Airyscan, and AI-assisted Airyscan micrographs, we found that AI-powered imaging achieved significantly higher resolution than the other two methods, capable of resolving structures less than 80 microns in width beyond the diffraction limit of light (**Fig. 4i**, right). Additionally, we revealed more detailed phenomena of hTPX2 in the nucleus during interphase and in the Prophase spindle during a cell cycle (**Extended Data Fig. 8**). These results underscore the potential of *de novo* Syn-phage in evolving nanobodies for captivating applications, such as SR immunoimaging of endogenous proteins inside living cells.

## Discussion

In summary, we introduced the “*de novo* microorganism” called Syn-phage that can be rationally designed, chemically synthesized, rescued in a host, and applied in attractive applications. It is intriguing to consider that the earliest life forms on Earth, as evidenced by fossilized microorganisms for example, might have been non-encapsulated due to their simplicity^27, 28^. In this context, Syn-phage exemplifies an unencapsulated form of artificial microorganism in which genetic material is physically linked to phenotypic proteins. Importantly, a Syn-phage is adaptable for the insertion of a nanobody gene, thereby enabling evolution of a nanobody, benefiting further applications. For instance, using F12-containing Syn-phage **v1.3** led to the creation of a G112S nanobody with fivefold increase in nanobody:antigen binding ratio and more than doubled binding affinities. Subsequent application of the G112S-EGFP chromobody allowed detailed visualization of endogenous hTPX2 patterns surpassing the diffraction limit of light through an AI-assisted Airyscan immunoimaging pipeline.

In the field of synthetic biology, innovative applications of microorganisms—such as directed evolution of biological molecules —have garnered great interest^29^. Our study focused on utilizing Syn-phage for the evolution of nanobodies, particularly those targeting intrinsically disordered antigens known for their reduced immunogenicity. By integrating the Syn-phage based evolution pipeline with the cost-effective naïve phage library screening method^30^, we were able to efficiently generate improved nanobodies (**Extended Data Fig. 9**). While phage surface display and other techniques have proven effective in enhancing nanobodies^31^, the Syn-phage based evolution approach stands out for its simplicity. It relies solely on a circular DNA as the essential reagent and eliminates any risk of laboratory phage contamination. Additionally, this method also circumvents potential issues associated with phage envelope assembly and surface antibody presentation^32^. Nevertheless, Syn-phages currently rely on acute environmental temperature fluctuation to “infect” a host with a comprised efficiency. Therefore, future research could center around designing Syn-phages that demonstrate positive-infectivity. Although this pursuit poses considerable challenges^33^, achieving success would open up a novel avenue for screening and generating artificial humanized antibodies entirely from scratch, showcasing substantial potential in biomedical applications^34^.

*De novo*-created artificial microorganisms have significant potentials across various fields as they can bypass the complexity of naturally existing microorganisms while strategically harnessing their beneficial characteristics^9^. Building upon the “*de novo* microorganism” methodology, future research could focus on developing alternative artificial microorganisms that parasitize Eukaryotic cells such as yeast, insect, or mammalian cells, with the aim of producing or enhancing innovative protein therapeutics and vaccines^35^. Furthermore, with notable progress in rational macromolecular assembly^36, 37^ and large DNA fragment synthesis^38, 39^, it will be even possible to create compartmentalized^40^ artificial *de novo* viruses or phages that are fully designed and synthesized by human with novel functions unparalleled by nature-existing ones.

## Methods

### Mammalian cell culture

HeLa cells (Cat# CL-0101) were obtained from Procell Life Science & Technology Co., Ltd. (Wuhan, P.R. China) while HeLa cells stably expressing mCherry-α-tubulin (Cat# CY016) was obtained from Inovogen Tech. Co., Ltd. (Chongqing, P.R. China). The cells lines were short tandem repeat (STR) identified and proven to be HIV-1, HBV, HCV, mycoplasma, and other microorganisms free before culturing. Other reagents such as full DMEM (Dulbecco’s modified Eagle’s medium) and PBS (phosphate buffered saline) were also confirmed to be mycoplasma free before usage. HeLa cell culture was maintained at 37°C under 5 % CO^2^ in high glucose (4.5 g·L^−1^) DMEM (HyClone, #SH30243.01) containing 4 mM L-glutamine and sodium pyruvate and supplemented with additional 10 % fetal bovine serum (FBS) (HyClone, Cat# SV30087.03), 1 % non-essential amino acid (NEAA, 100×), and 1 % penicillin-streptomycin (100×). Trypsin-EDTA (HyClone, #SH30042.01) and PBS (HyClone, #SH30256.01) were used in subculturing. HeLa cells were subcultivated in a ratio of 1:5∼10. HeLa cells stably expressing mCherry-α-tubulin were maintained at 37°C under 5 % CO^2^ in minimum essential medium (MEM) (PriCella, Cat# PM150414) containing 4 mM L-glutamine and sodium pyruvate and supplemented with additional 10 % fetal bovine serum (FBS) (HyClone, Cat# SV30087.03), 1 % non-essential amino acid (NEAA, 100×), 1 % penicillin-streptomycin (100×), and 0.1 % puromycin (Shanghai MaokangBio, Cat# MS0011-100MG). Trypsin-EDTA (HyClone, Cat# SH30042.01) and PBS (HyClone, Cat# SH30256.01) were used in subculturing. mCherry-α-tubulin HeLa cells were subcultivated in a ratio of 1:2∼3.

### Plasmid construction

Plasmid vectors, such as pTXB1, pET28a(+), EGFP-C1 and EGFP-N1 were obtained from commercial vendors. These parental vectors may be further engineered, such as introducing a His^6^- or His^8^-affinity tag, insertion of a TEV (ENLYFQ↓G) protease cleavage site, alternation of restriction cleavage sits, or replacing EGFP by other fluorescent proteins, e.g. mCherry, etc. to give modified versions of the parental vector for cloning. Subcloning, Gibson cloning, or modified Gibson cloning methods were employed to construct the desired plasmids. For subcloning, fragments of interest were directly cut from the parent plasmid using appropriate restriction enzymes, or amplified by PCR from plasmids containing the desired genes using hyPerFUsion high-fidelity polymerase (APExBIO, #1032,), gel purified, digested with restriction enzymes and purified again. The gene fragments were ligated into appropriate vectors using T4 DNA ligase. Multiple fragments were assembled by stepwise subcloning or one-step multi-fragment Gibson cloning. Custom gene synthesis was received from Ruibiotech (Beijing, P.R.China), General Biol (Chuzhou, P.R. China), or Comate Bioscience (Changchun, P.R. China).

### Transfection

Transient transfection was typically performed in an 8-well (Cat# 155409) Lab-Tek®II imaging chamber from Thermo Scientific using Lipo8000™ transfection reagent from Beyotime Biotechnology (Cat# C0533). Typically, 0.25 μg DNA was dissolved in 12.5 µl gibco opti-MEM (Life technologies, Cat# 31985-062) and then 0.4 μl Lipo8000™ transfection reagent was added and mixed via gentle pipetting. Then this mixture was added into an imaging chamber well seeded with 2.5×10^4^ cells that were already adhesively attached on the bottom in 250 μl full DMEM. The cells were maintained under 5% CO^2^ at 37 °C for around 2 h. Then the medium was replaced by warm full DMEM and the cells were further incubated under 5% CO^2^ at 37 °C for over 20 h. For co-transfection of more than one plasmid, the quantity of DNA used in this protocol implies the total amount of plasmids.

### Confocal microscopy and AI-assisted Airyscan imaging

Live cells were imaged in phenol red free Dulbecco’s Modified Eagle Medium (Life Technologies, Cat# 21063-29) supplemented with additional 10 % FBS, 1 % sodium pyruvate, 1 % NEAA, 1 % penicillin-streptomycin and 15 mM HEPES-Na at 37° under 5 % CO2. Microscopy was performed using Nikon A1 ECLIPSE T*i*2 inverted confocal microscope. The microscope was equipped with four lasers (405 nm, 488 nm, 561 nm, and 640 nm), allowing the acquisition of confocal fluorescence data for four different excitation wavelengths. For detection of blue (excited by 405 nm laser) or far red (excited by 640 nm) fluorescence signal, PMT (photomultiplier tube) detectors were used; while for the detection of green (excited by 488 nm) or red (excited by 561 nm) fluorescence signal, the more advanced GaAsP detector with even higher sensitivity will be used. Most images presented in this article were acquired with a 60× oil objective lens (APO 60×/1.40 oil) having a numerical aperture of 1.4. In most cases, the basic imaging setup were configured with typical parameters set as follows: Scan speed 0.5, number of averaging 4, scan line mode in one-way scan direction.

Alternatively, microscopy was performed using Zeiss LSM 880 inverted confocal laser scanning microscope equipped with an Airyscan super-resolution module. This microscope was equipped with a 405 nm laser diode, 458/ 488/ 514/ nm argon laser, and 543 HeNe laser and 594 HeNe laser for excitation of different fluorophores. Zeiss Plan-APOCHROMAT 63×/1.4 oil DIC objective was used as the primary objective. Confocal images were typically acquired in 12-bit depth at 512×512 resolution. In most cases, the basic imaging setup parameters were configured applying the “Smart-Setup” function with typical parameters set as follows: scan speed 8, pixel dwell time 1.54 μs, number of averaging 4, line mode in one-direction scanning, and pinhole 89.9 μm. Where appropriate, edges of live cells recorded in microscopy were sketched as white lines for better clarity in this study. To acquire AI-assisted high-resolution micrographs, Airyscan module was used by selecting the ChA channel and acquiring images at 1024×1024 resolution. Subsequently, the obtained images were opened by PixInsight (https://pixinsight.com) and processed by applying BlurXTerminator (drag the triangular icon in this process onto the opened image) to create AI-assisted Airyscan super-resolution images.

### Immunofluorescence (IF) staining

HeLa cells were washed three times by PBS, fixed using 4 % paraformaldehyde for 20 min, washed three times by PBS, permeabilized using 0.5 % Triton X-100 in PBS for 20 min, washed three times by PBS, blocked with 5 % BSA in PBS at room temperature for 30 min. Blocking solution was removed and primary antibody against hTPX2 (Cat# R27376, ZENBIO) diluted 1:100 in 0.5 % BSA/ PBS was added and cells were incubated at 4 °C overnight. Next day, primary antibody solution was removed and then cells were incubated with Alexa Fluor 488 labeled secondary antibody (Cat# 550037, ZENBIO, 1:100 diluted in 5 % BSA/ PBS) at room temperature for 1 h. Secondary antibody was removed and cells were washed three times by PBS and imaged under a confocal microscope.

### Protein expression and purification

pET28a(+) or modified pET28a(+) vectors were used in most protein expressions while pTXB1 vector was used to express intein-tag fused proteins such as nanobody-intein chimeras for expressed protein ligation (EPL). These plasmids for protein expression were first transformed into *E. coli* Rosetta 2a cells and the transformants were selected on ampicillin (125 mg·L^-1^) or kanamycin (50 mg·L^-1^) agar plates depending on the antibiotic resistance of the plasmids. A single colony was used to inoculate 50-100 ml of LB medium containing 125 mg·L^-1^ ampicillin or 50 mg·L^-1^ kanamycin and shaken at 240 rpm for 8-10 hours or overnight at 37 °C. 30-50 ml of the preculture was used to further inoculate ∼1.8 L fresh LB medium containing 125 mg·L^-1^ ampicillin or 50 mg·L^-1^ kanamycin, and additional chloramphenicol (33 mg·L^-1^). The absorbance at 600 nm (OD600) of the inoculated culture should be controlled between 0.05 to 0.1 in this inoculation step. Then the culture was shaken at 180 rpm at 37°C for a few hours (typically 2-3 h) until OD 600 reached 0.5-0.6. Then 0.5 ml isopropyl β-D-thiogalactoside (IPTG) stock solution (1M) was added (final ∼0.27 mM) to induce protein expression at 30 °C or 16 °C overnight. Sometimes protein expression time and temperature needed to be optimized in order to achieve an optimal expression for some particular proteins.

Later, cells were harvested by centrifugation at 13881× g, at 4 °C for 15 min and washed once with PBS (4149× g, 10 min). The bacterial pellet was resuspended in lysis buffer (pH 8.0, PBS supplemented with additional 0.5 M NaCl, 3 % glycerol, w/o 3 mM β-mercaptoethanol (BME), and 1 mM phenylmethylsulfonyl fluoride (PMSF). For relatively smaller volumes of bacterial cell suspensions (< 40 ml), bacterial cells were typically lysed via ultra-sonification at 80 W for 30 min or 60 W for 45 min (1 s sonification followed by 3 s interval) on ice. For batch processing or larger volumes of cell suspensions, cells were typically lysed using ultra-high-pressure homogenizer cooled by a bench chiller for 2-3 cycles under 800-900 bar at 4 °C. The lysate was cleared by high-speed centrifugation (74766× g, 45 min, 4 °C) and the supernatant was loaded onto a gravity Ni-NTA column (2-5 ml Ni-charged resin FF from GenScript, Cat# L00666-25). The Ni-NTA column was washed and then the His-tag fused protein was eluted using step-gradient of imidazole (50, 100, …, until 500 mM) solutions. Size exclusion chromatography (SEC) or ionic exchange may be further applied using GE ÄKTA Pure if additional purifications are necessary. The obtained proteins were typically concentrated, buffer exchanged in buffer A (pH 8.0 PBS, 0.5 M NaCl, 3% glycerol, w/o 3 mM BME), aliquoted, snap frozen in liquid nitrogen, and stored under -80 °C.

### Generation of the lead nanobodies from naïve libraries via M13 phage display

The lead nanobody C3 against the intrinsically disordered protein BuGZ was obtained using BuGZ as the antigen to screen another naïve phage library. A modified pCANTAB5e phagemid vector was used and panning and ELISA selection were performed in Atagenix company. BuGZ was expressed and purified with a qualified purity, stored at -80 °C as snap frozen aliquots, and thaw at 4 °C overnight just before use. 40 μg BuGZ in 1 ml PBST or 5% milk-PBST (control) was used to coat Nunc immune tube (Cat# 444202, Thermo Scientific) at 4 °C overnight. Panning was performed according to the standard operation protocol (SOP) of the company, which produced a sub-phage library ready for ELISA selection. After three rounds of panning and several ELISA detections, total eleven positive sequences were obtained and the G3 nanobody was identified to be a strong binder against BuGZ. The lead nanobodies (G6 and F12) against hTPX2 were obtained via screening another naïve phage library using a similar process. For this, pADL10b phagemid was used and panning and ELISA selection were performed in AlpaLife company.

### Isothermal titration calorimetry (ITC)

ITC was performed using MicroCal ITC200 from GE Malvern. Nanobody and the antigen were all dissolved in freshly prepared pH 7.2 PBS buffer supplied with 0.5 M NaCl, 3 % glycerol, and additional 0.4 mM TCEP as reductant. 400 μl of antigen at specified concentrations was loaded into the sample cell. Nanobody as specified concentration was loaded in the syringe and then injected in 2.0 μl × 18 portions to the sample cell at 25 °C with 3 min interval between each injection; only for the first injection, 0.8 μl nanobody was used followed by 2.5 min interval. The titration data was processed using the default Origin software. The following information describes the concentrations used for each ITC measurement: G6 at 300 μM titrates hTPX2 at 69 μM; KV at 600 μM titrates hTPX2 at 30 μM; F12 at 50 μM titrates hTPX2 at 20 μM; G112S at 150 μM titrates hTPX2 at 20 μM; C3 at 400 μM titrates BuGZ at 36 μM; W102A at 300 μM titrates BuGZ at 35 μM.

### Förster resonance energy transfer (FRET)

EGFP and mScarlet are a FRET pair due to the spectra overlap between the emission spectrum and absorption spectrum of the two. The Molecular Device (MD) *SpectraMax i3x* spectrometer was used in FRET measurement. 100 μl or 200 μl solution of a mixture of donor and acceptor fluorophores were added into each well in the black 96-well plate. The excitation wavelength was set at 470 nm and the fluorescent spectra were recorded from 490-750 nm.

### Identification of key residues in CDR regions for evolution

First, the CDR regions of a nanobody were predicted using *The International Immunogenetics Information System* (IMGT, web: https://www.imgt.org/) under IMGT/DomainGapAlign. The CDR region is the position where evolution is going to be performed. Meanwhile, the SWISS-MODEL (web: https://swissmodel.expasy.org/) online prediction tool can be used to predict the 3D structure of the nanobody and the respective PDB file can also be generated. With this PDB file in hand, alternative softwares, such as the freely accessible PyMOL (web: https://pymol.org/2/) can be used to facilitate visualization of the predicted 3D structures and identification of the CDR and non-CDR regions of a nanobody.

### Syn-phage based directed evolution protocol

#### General

Syn-phage-based nanobody evolution consists three major steps, **i**) gene diversification, **ii**) rescue and screening, **iii**) characterization of nanobodies. The following section describes the evolution of C3 nanobody which is a representative protocol involving single-site saturated mutagenesis covering the entire CDR3 region. All steps are belonging to standard molecular biology or biochemistry procedures.

#### Step I: Gene diversification

Eleven PCR reactions were performed using the forward primer and each of eleven single site-saturated mutagenesis reverse primer to produce the 5’-end diversified C3 nanobody gene fragment; meanwhile, another eleven PCR reactions were performed using each of eleven single site-saturated mutagenesis forward primer and the reverse primer to produce the 3’-end diversified nanobody gene fragment (see Supplementary Information for primer sequences). Also, the circular DNA of the parental Syn-phage **v1.0** was cut by NheI and NdeI to produce the vector template. Therefore, total eleven one-pot ligation reactions catalyzed by recombinase were performed to simultaneously insert the N-half and C-half of the nanobody gene fragments into the Syn-phage **v1.0** gene template, creating circular Syn-phage **v1.2** gene with single site-saturated mutagenesis at the entire CDR3 region of C3 nanobody. The ligation products were transformed into Rosetta 2a chemical competent cells in eleven individual 1.5 ml vials. Each transformant suspension (∼25 μl) was first mixed with 1 ml of antibiotic-free LB medium shaking at 37 °C for 1h before inoculating 10 ml LB medium containing 50 mg·ml^-1^ kanamycin in a test tube. The next day, 4 ml culture in each test tube was subjected to plasmid mini-prep, and the obtained eleven groups of diversified Syn-phage genes were subjected to Sanger sequencing to confirm that successful creation of mutations at the CDR3 region. At the sample time, 0.5 ml culture in each test tube was mixed with 0.5 ml 50 % glycerol and stored at -80 for future use.

#### Step II: Rescue and selection

The remained 5.5 ml bacterial culture in each of the eleven test tubes were combined, suspended in 1.8 L LB (50 mg·ml^-1^ kanamycin & 33 mg·ml^-1^ chloramphenicol), and subjected to standard bacterial protein expression (OD600 0.6, then 25 °C overnight). The Syn-phage **v1.2** mutant library was first isolated using Ni-NTA affinity column chromatography, purified by SEC, and screened over BuGZ-coated beads. We used NHS Focurose 4FF bead (Cat# HQ030302001, HUIYAN Bio, Wuhan, China) for the preparation of BuGZ-coated bead following the manufacture’s protocol. NHS Focurose 4 FF is a kind of NHS bead that can covalently immobilize proteins on its surface via the reaction between lysine-reactive NHS group on the beads and the lysine side chains of a protein. For a typical screening process, 150 μl purified Syn-phage **v1.2** library was mixed 50 μl BuGZ-coated beads, incubated at 4 °C for 1 h, and finally washed three times by PBST. Therefore, these beads will bind Syn-phages that carry evolved C3 nanobodies with enhanced binding affinity.

#### Step III: Characterization of nanobodies

The evolved nanobody genes were amplified by performing PCR reactions of the above screened beads using the forward and reverse primers of the lead nanobody. The nanobody amplicon library was inserted into pCMV_EGFP-mito vector to give pCMV_C3(mutant)-EGFP-mito plasmids by subcloning using Rosetta 2a competent cells. The next day, forty-eight Rosetta 2a colonies on a Kan agar plate were picked and subjected to Sanger sequencing and those mutations show two or more repeating frequency potentially suggest that beneficial mutations occurred. According to sequencing results, total eight nanobodies were identified, which were W102A, D112A, R104K, N105M, N105T, K108P, R104D, and N105V. These C3(mutant)-containing plasmids were subjected to live-cell colocalization assay revealing that W102A, D112A and R104K nanobody mutant show higher colocalization degree with BuGZ. Finally, the three nanobodies are further characterized, for example via ITC to determine their binding affinities and other binding thermodynamic parameters.

### Quantification and Statistical Analysis

#### Image analysis

Microscopic images were analyzed and processed with ImageJ/Fiji and prepared for presentation using Microsoft Office PowerPoint. Image manipulations were restricted to adjustment of brightness level (i.e. linear stretch), background subtraction, cropping, rotating, scaling, and false color-coding using Look-Up Tables (LUT). Pearson’s correlation coefficient (PCC) as well as Manders correlation coefficient (MCC) was employed for colocalization analysis using the “Manders_Coefficients.class” plugin for ImageJ/Fiji. Usually, there is only a single cell within an imaging field; otherwise, a single cell was first selected using the polygon selections tool and then remove extra cells (Edit/Clear outside) prior to PCC calculation. The images were converted to 8-bit depth (Image/Type/8-bit) prior to analysis, and typically no less than five cells were analyzed for each PCC or MCC.

#### Statistics and reproducibility

All microscopic imaging experiments were representative of at least three independent repeats if not otherwise stated; representative SDS-PAGE images were from at least three independent repeats with similar results; representative confocal microscopic images were from at least five cells with similar results. No randomization nor blinding was used in this study. Origin and Microsoft Excel were used for plotting, data fitting, graphing and statistical analysis. All box plots show mean (square), median (bisecting line), bounds of box (75^th^ to 25^th^ percentiles), outlier range with 1.5 coefficient (whiskers), and minimum and maximum data points (lower/ upper whiskers). Student’s *t*-tests were used to compare two experimental conditions. Unless otherwise specified, one-sided unpaired *t*-tests were performed, as for example cells expressing different nanobody mutants. When necessary, stars were used to denote *P*-values for indicated statistical tests (*: *P*<0.05; **: *P*<0.01; ***: *P*<0.001; ****: *P*<0.0001). Exact *P*-values were indicated for critical experiments.

## Code availability

“Manders_Coefficients.class” plugin and how to use it in ImageJ/Fiji for the calculation of PCC values were provided as a supplement in a previously published paper (PMID: 36964170).

## Acknowledgement

We thank the Core Facilities at HCLS and School of Life Science and Technology in Harbin Institute of Technology. We thank Prof. Panpan Wang and Zhilin Zhang for performing atomic force microscopy experiments and valuable discussions; we thank Siyang Li and Qinqin Jiang for technical assistance, and Chengjian Zhou for fixed cell immunofluorescence imaging experiments. This work was supported by the funding from Harbin Institute of Technology, Overseas Outstanding Young Talents Program of China, National Natural Science Foundation of China (grant No. 32071410 to X.C.), Natural Science Foundation of Heilongjiang Province of China (grant No. YQ2022B004 to X.C.).

## Author contributions

X.C. conceived and supervised the project. X.C., D.W, and H.H. performed formal analysis and contributed to interpretation. X.C., D.W. and H.H. prepared all the figures. X.C. wrote the paper and D.W. and H.H. contributed to proof-reading. D.W. and H.H. performed most of the experiments with contributions from R.X. and S.X. D.W. performed all ITC experiments including the expression and purification of the corresponding antigens and antibodies. D.W. performed all immunoimaging and AI-powered Airyscan imaging. H.H. achieved expression and purification of Syn-phages and characterization of them including SDS-PAGE, Western-blot analysis, and MS^2^ mass fingerprinting.

